# A simple egg marking method for polygynous fishes

**DOI:** 10.1101/502179

**Authors:** Tessa K. Solomon-Lane, Matthew S. Grober

**Affiliations:** Department of Integrative Biology, 1 University Station #C0930, University of Texas at Austin, Austin, TX 78712;.; Department of Biology, 50 Decatur Ave, Georgia State University, Atlanta, GA 30302; Neuroscience Institute, 50 Decatur Ave, Georgia State University, Atlanta, GA 30302

**Keywords:** egg, dye, mark, external fertilization, fish, food coloring, *Lythrypnus dalli*, reproductive success

## Abstract

Fitness is the ultimate measure of organismal function; however, technical challenges can limit researchers’ ability to quantify fitness. We developed a simple and inexpensive method of marking eggs inside of the ovary of a nesting fish, the bluebanded goby (*Lythrypnus dalli*), in order to quantify female reproductive success. Multiple females in a harem of *L. dalli* can lay eggs inside of the male’s nest within a short period of time, making the timing of egg laying or developmental stage of the eggs insufficient to identify which female laid which clutch of eggs. After injecting small volumes of food coloring into the ovaries, each female’s eggs can be identified throughout development by the color of the yolk. We make preliminary observations about the efficacy of different colors (red, yellow, green, blue) for use in *L. dalli*, including effects on female survival, social behavior, and the enzyme immunoassays used to quantify hormones. While rigorous validations should be conducted for each species and experimental context, this method of marking eggs has the potential to be broadly useful for directly estimating fitness in species with external fertilization, as well as research in reproductive biology, development, behavioral ecology, and evolution.

## INTRODUCTION

Quantifying fitness is fundamentally important to testing the adaptive function of traits in the study of evolution and its underlying mechanisms. Reproductive success is one of the best approximations of fitness (reviewed in Pradhan et al., 2015), and it is feasible to quantify the number of offspring produced by an individual in a diversity of species in the field (e.g., Buston, 2004; Griffith et al., 2008; Silk et al., 2009) and in the laboratory/artificial habitats (e.g., O’Rourke and Mendelson, 2014; Sih et al., 2014; White et al., 2010). Here, we report on a simple and inexpensive method of marking fish eggs with food coloring in order to quantify reproductive success for multiple females simultaneously in a harem of bluebanded gobies (*Lythrypnus dalli*). This highly social, sex changing fish is a useful study species because females lay large numbers of demersal eggs frequently (Solomon-Lane et al., 2015), males readily parent (Pradhan et al., 2014), and the hatched larvae can be reared in the laboratory (Archambeault et al., 2016). Quantifying male reproductive success has already provided important insight into social behavior and social network structure as targets for natural selection (Solomon-Lane et al., 2015).

Across species, parentage cannot be assumed on the basis of which nest offspring develop in or the identity of the individuals providing parental care. For example, extra pair copulations in birds (Griffith et al., 2008) and pirate (e.g., Tatarenkov et al., 2006) and sneaker males (e.g., Svensson and Kvarnemo, 2007) in fish can result in parents caring for unrelated offspring. For gobies, one of the largest families of advanced fishes, multiple females can lay eggs in the same nest within short succession (Tamada, 2008). Visual assessment of developmental stage or the timing of egg laying are not sufficient to identify which female contributed which clutch of eggs (Takahashi and Ohara, 2006). Our goal was to develop a method of marking eggs in order to rapidly and visually identify the mother while having minimal/no impact on female health, social behavior, or reproductive behavior and biology. Previous studies have used intraperitoneal injections of various dyes, the most effective of which for gobies was brilliant blue FCF (Okuda et al., 2002). We expand on this work by testing multiple colors of dye that are commercially available as food coloring. Food coloring is inexpensive, multiple colors can be delivered in the same vehicle (see Okuda et al., 2002), and, to our knowledge, the dyes do not independently impact reproductive success (e.g., β-carotene, Okuda et al., 2002; Olson and Owens, 1998).

We used two small injections of dye, one into each ovary, to mark eggs (Fig 1a, b). The dye incorporates into the egg yolk and persists through embryonic development (Fig 1c). This manipulation was used as a part of a study investigating connections among social behavior, reproduction, and hormones, and we share preliminary observations about effects on *L. dalli* health, social interactions, egg laying, and the enzyme immunoassays used to quantify hormones. Beyond this proof of principle, direct validations of this method for other contexts and/or other species should be conducted to identify and assess possible side effects or unintended consequences. This simple method of dying eggs can be broadly useful for estimating female fitness in species with external fertilization.

**Figure 1:**
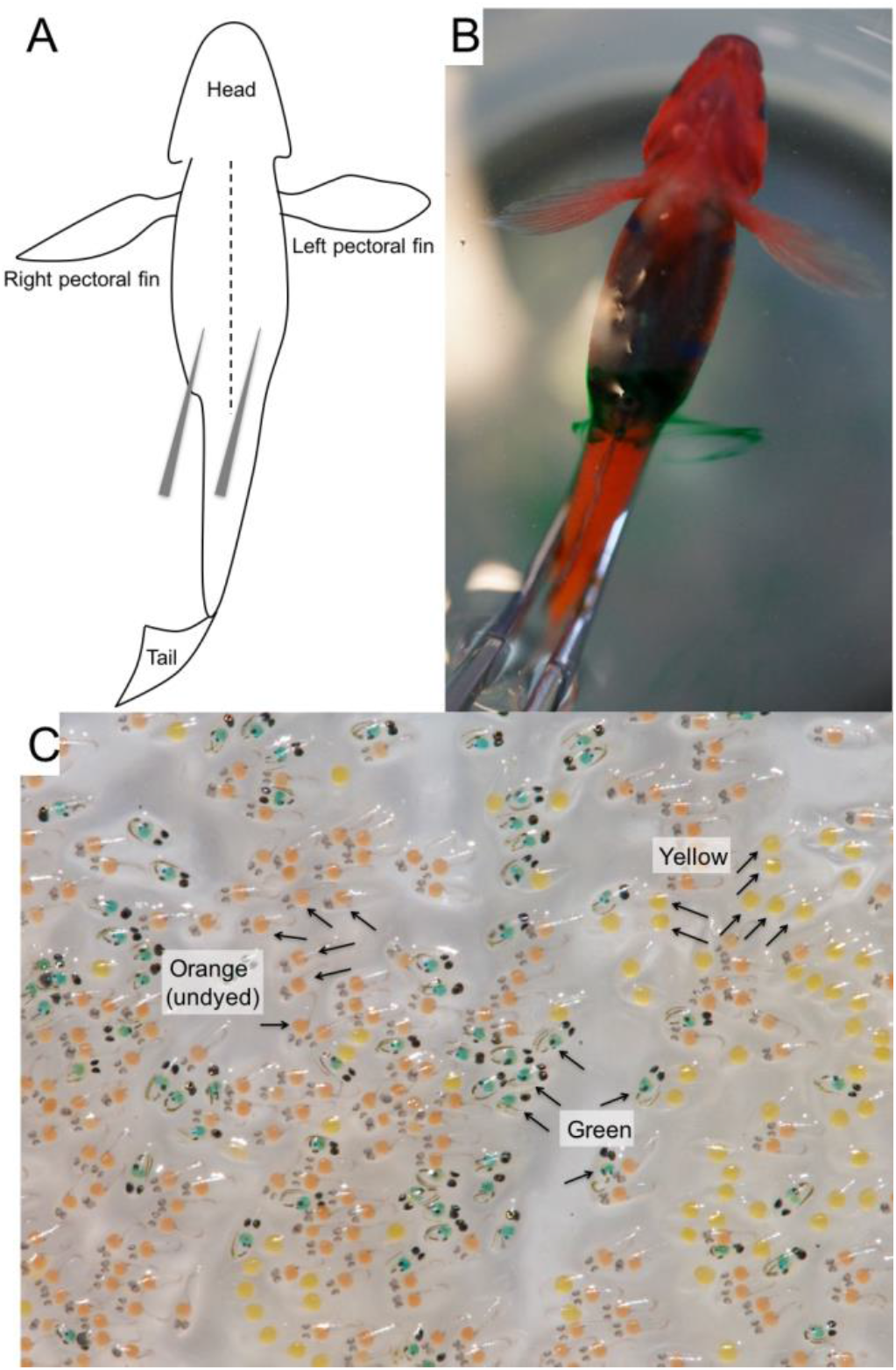
Injections of food coloring into the ovary to mark eggs. **A)** Schematic drawing (ventral view) of *L. dalli* with injection sites indicated. **B)** Digital image (ventral view) of an anesthetized gravid female immediately post-injection of green dye. A small trail of dye is leaving one injection site. **C)** Digital image of multiple egg clutches laid in the laboratory on a sheet of acetate. The eggs with a yellow yolk were newly laid by a female injected with yellow dye. The eggs with a green yolk, developed eyes, and a visible tail were laid by a female injected with green dye. The eggs with an orange yolk are developmentally intermediate and were laid by an uninjected female.

## MATERIALS AND METHODS

### Social group formation and dye injection

We collected *L. dalli* (23.5–47.6 mm standard length) from reefs offshore of Catalina Island, California during the reproductive season (California Fish and Game permit SC-11879) using hand nets while SCUBA diving. The fish were housed at the Wrigley Institute for Environmental Studies (Catalina Island, University of Southern California). One day after collection, we formed social groups of 4 fish: 1 large, dominant male and 3 females of varying sizes (n=65). Behavior and reproduction were analyzed in social groups with no mortality (n=51). To form groups with fish of specific sizes and sex ratios, fish were briefly anesthetized in tricaine methanesulfonate (MS-222; 500 mg/L salt water). To dye the eggs, we injected a small volume (∼0.03–0.05 mL) of undiluted blue, yellow, or green food coloring (Dec-A-Cake icing tint) into each ovary using a 28.5-gauge insulin syringe (BD Lo-Dose) (Fig 1a). Each female in a group was dyed a different color, and color was balanced across statuses. We did not include a no-dye control injection group in this study (see discussion). Our pilot experiments suggest that the stage of egg development in the ovary can impact the effectiveness of the dye. For *L. dalli*, mid-cycle injections were the most effective. Females injected immediately after laying often had clutches that were incompletely or only faintly dyed, similar to intraperitoneal (rather than ovarian) injections.

### Quantifying reproduction and behavior

Males were provided with a PVC nest tube (3 inch length, 1 inch diameter) lined with acetate, on which females lay demersal eggs. We performed egg checks from the outside of the tank every 30 minutes from 6 am to 8 pm, and if present, we removed the acetate, took a digital picture, and returned the eggs within minutes (as in Solomon-Lane et al., 2014). We used ImageJ (Schneider et al., 2012) to count the number of eggs laid.

Dye could affect social behavior through physiological mechanisms or because it temporarily (∼24 hours) colors the entire fish, internally and externally (Fig. 1c). We conducted 4 10-minute behavioral observations in the first ∼52 hours after social groups were formed, including 1 min, 3 hours, ∼24 hours, and ∼52 hours after the fish were introduced into their group. We recorded social interactions among all members of the group, including approaches, when one fish swims directly towards another fish within 2 body lengths, and displacements, a response to an approach in which the approached fish swims away. Displacements are a measure of aggression, and being displaced is a signal of submission by subordinates (Rodgers et al., 2007). Behaviors summed over the 4 observation periods are presented as behaviors per min. We tested for effects of dye color on the behaviors expressed by the dyed fish (approaches, displacements), as well as how group members interacted with that fish (approaches to the focal female, submissions by/displacements of the focal female). Because the effects could be status-specific (Solomon-Lane et al., 2014), we also included social status in our analyses.

### Enzyme immunoassays

Collecting water-borne hormones is a non-invasive method of quantifying systemic steroid hormones (Kidd et al., 2010). Water collected from injected fish can contain visible quantities of dye; therefore, it was critical to determine whether dye affected the enzyme immunoassays (Cayman Chemical). To test the effect of the green, yellow, and blue dyes on the cortisol, 11-ketotestosterone (a potent fish androgen), and 17β-estradiol enzyme immunoassays, we completed two standard curves per assay. We added 1μl of dye to each well of one standard curve (e.g., cortisol: yellow in standards 2–3; blue in standards 4–5; and green in standards 6–8). 1μl of ultrapure water was added to wells of the control curve. The assay was then completed according to the supplied instructions, and the plates were read 105 min following development for cortisol and 11-ketotestosterone and after 135 min for 17β-estradiol. We present the equation and r-squared value for the dyed and control standard curves.

### Data analysis

Statistical analyses were conducted using R Studio (version 1.0.143). Results were considered significant at the p<0.05 level. The box of the box and whisker plots show the median and the first and third quartiles. The whiskers extend to the largest and smallest observations within or equal to 1.5 times the interquartile range. We used mixed factorial ANOVAs to identify differences in behavior (approaches, displacements, approaches to females, submissions) among females injected with different dye colors (between-subjects factor), differences among females of different statuses (within-subjects factor), or status-by-color interactions. Tukey’s HSD tests were used for *post hoc* analysis of significant results. Chi-squared tests were used to analyze differences across dye colors in mortality and egg laying. A Mann-Whitney U test was used to compare the number of eggs laid by yellow- and green-injected females. Clutches of blue eggs were excluded due to insufficient sample size (n=3). Statistical comparisons of the slopes of dyed and control standard curves were compared by hand as in Fischer et al., 2014 (Zar, 1999).

## RESULTS

### Dye colors and mortality

Red, yellow, blue, and green dyes are included in standard food coloring packages. Red dye (no. 40) was not used because the fish did not survive the pilot injections. We formed 65 social groups of 1 male and 3 females of different social statuses (alpha, beta, gamma). Each female in the group was injected with a different dye color (yellow, green, or blue), and dye color was balanced across social statuses. Mortality was very low overall (179 of 195 females survived; 8.2% mortality), and there were no differences in survival across dye color (χ^2^=0.54, n=195, d.f.=2, p=0.76).

### Dye color does not affect social behavior

There were no differences in rates of approaching among dye colors (*F*_2,144_=2.15, p=0.12), but there was a trend for differences in rates of approaching across social statuses (*F*_2,144_=2.96, p=0.055). There was no interaction effect (*F*_4,144_=0.62, p=0.65). For displacements, there were no differences among dye colors (*F*_2,144_=0.84, p=0.44), but social statuses differed significantly (*F*_2,144_=7.75, p=0.00064). There was no interaction effect (*F*_4,144_=0.87, p=0.48). *Post hoc* tests showed that alphas (p=0.00062) and betas (p=0.014) displaced other fish significantly more than gammas. There were no differences between alphas and betas (p=0.60) (Fig 2b).

**Figure 2:**
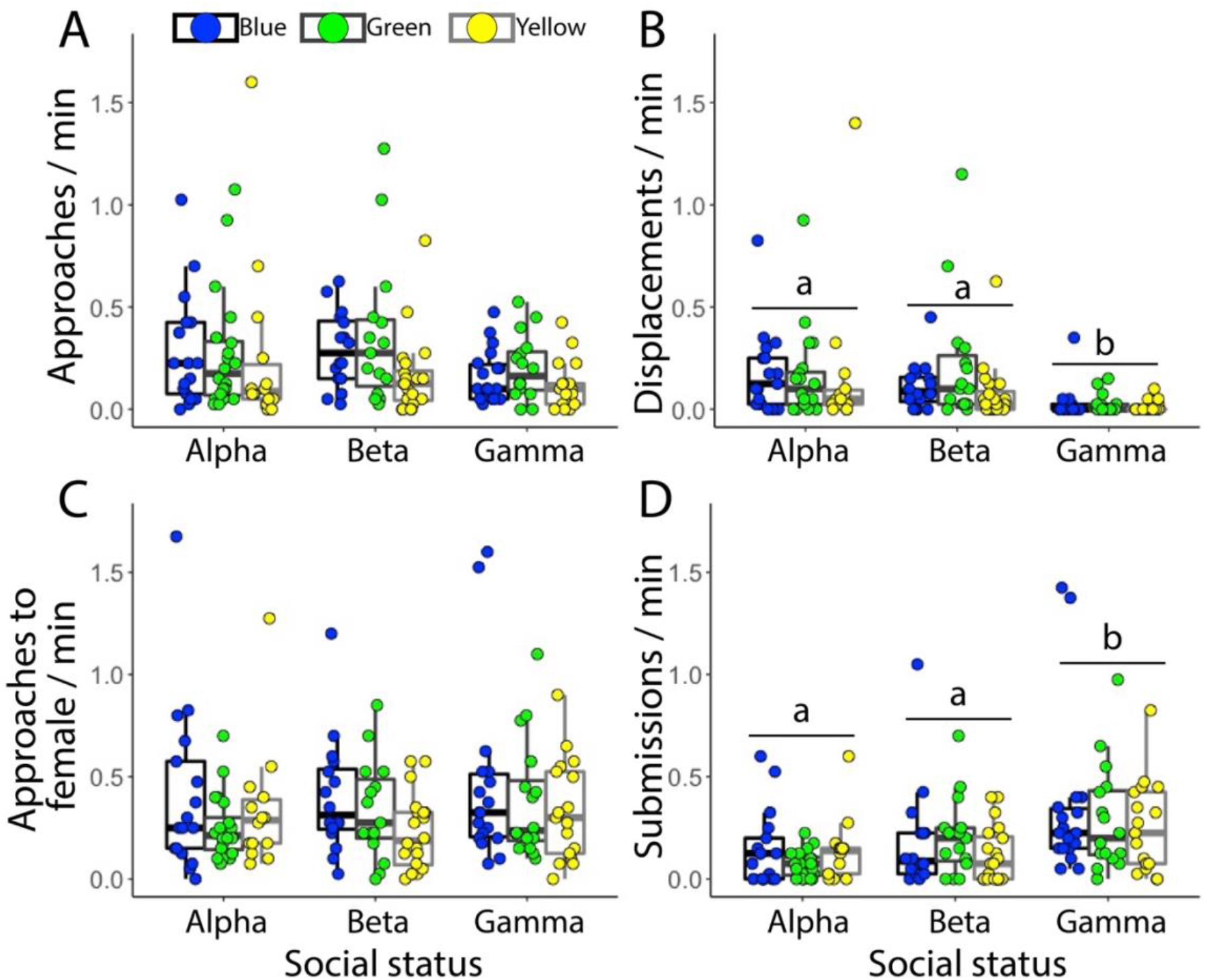
Social behavior across dye color and social status. **A)** Mean (±SEM) approaches and **B)** displacements by alpha, beta, and gamma females injected with yellow, green, and blue dye. **C)** Mean (±SEM) approaches directed to dyed females by other group members. **D)** Mean (±SEM) submissions by dyed females. Yellow alpha (n=14); yellow beta (n=20); yellow gamma (n=17); green alpha (n=20); green beta (n=15); green gamma (n=16); blue alpha (n=17); blue beta (n=16); blue gamma (n=18). Different letters indicate significant status differences (p<0.05).

There was a trend for other members of the social group to approach the focal fish differentially based on dye color (*F*_2,144_=2.73, p=0.068) but not status (*F*_2,144_=1.04, p=0.35). There was also no interaction (*F*_4,144_=0.50, p=0.74) (Fig 2c). Finally, there were no differences in rates of submission (i.e., focal fish displaced by others) among dye colors (*F*_2,144_=1.04, p=0.34), but there were significant differences across social status (*F*_2,144_=9.04, p=0.0002). There was no interaction effect (*F*_4,144_=0.59, p=0.67). *Post hoc* tests showed that gammas submitted significantly more than alphas (p=0.00017) and betas (p=0.0087). There were no differences between alphas and betas (p=0.50) (Fig 2d).

### Dye color affects egg laying

Dye color had a significant effect on the proportion of females that laid eggs (χ^2^=6.80, N=153, d.f.=2, p=0.033). Blue-injected females were significantly less likely to lay than either green (χ^2^=6.04, n=102, d.f.=1, p=0.014) or yellow females (χ^2^=6.04, n=102, d.f.=1, p=0.014). There were no differences between green- and yellow-injected females (χ^2^=0, n=102, d.f.=1, p=1.00). Of the females that laid eggs, there was no significant difference in the number of eggs laid by yellow- (average 1200.3 ± 248.3 eggs) and green-injected (average 991.7 ± 216.9 eggs) females (p=0.93) (average undyed in Solomon-Lane et al., 2015: 892.5 ± 62.8)

### Dye color does not affect enzyme immunoassays

We found that the slopes of standard curves spiked with yellow, green, and blue dye did not differ significantly from the control standard curves for cortisol (t_12_=0.034, p=0.97; control: y=–1.43ln(x)+6.65, r^2^=0.99; dye: y=–1.16ln(x)+5.06, r^2^=0.99), 11-ketotestosterone (t_12_=0.30, p=0.98; control: y=–1.22ln(x)+1.58, r^2^=0.99; dye: y=–1.10ln(x)+1.10, r^2^=0.99), or 17β-estradiol (t_9_=0.31, p=0.76; control: y=–0.96ln(x)+3.83, r^2^=0.98; dye: y=–0.94ln(x)+3.39, r^2^=0.99).

## DISCUSSION

In this proof of principle, we demonstrate that food coloring can be used to effectively mark live *L. dalli* eggs prior to egg laying. This method is simple and inexpensive, and due to culinary demand, a wide variety of colors are commercially available as liquid or powder. By counting the number of colored eggs laid inside of a male *L. dalli*’s nest, we can quantify individual female reproductive success in a social group with multiple females. The ability to estimate reproductive success for all members of a group will allow us to directly investigate the evolution of social and reproductive behavior (e.g., Solomon-Lane et al., 2015), as well as the underlying neuroendocrine mechanisms (e.g., Pradhan et al., 2014).

Our preliminary analyses suggest that green and yellow dyes will be the most useful for marking *L. dalli* eggs, although additional colors that have yet to be tested may work equally well. Females injected with green and yellow had high survival, laid eggs, and interacted socially. Comparisons to a no-dye control injection group will ultimately be necessary for identifying any specific effects on health, social behavior, male parenting, as well as egg laying, development, and hatching success. Based on personal observation, males parent dyed eggs, which go on to develop (Fig 1c) and hatch. Green, yellow, and blue dyes also had no effect on the standard curves for cortisol, 11-ketotestosterone, and 17β-estradiol, suggesting these dyes can be used in experiments that quantify hormone concentrations in water, blood, or tissue using enzyme immunoassays. This method of quantifying neuroendocrine function is very common and can be used to understand the regulation of behavior and reproduction (e.g., Pradhan et al., 2014).

For species like *L. dalli*, in which females lay multiple clutches during the reproductive season, an important limitation of this method is the need to re-inject a female each time she lays eggs in order for her next clutch to be marked. Females that lay more frequently will receive more injections, and groups with higher rates of reproduction will be disturbed more frequently. The potential negative impacts of these stressors on social and reproductive dynamics remain to be tested, although *L. dalli* tolerated multiple injections well. There may also be methods for releasing dye over time in order to avoid repeated injections. One possible solution is implants made by mixing gelatin powder with food coloring. A small piece of the gelatinous material will release dye into phosphate buffer for more than 3 weeks, and the mixture can be injected while warm into anesthetized females so that it gels *in situ* in the abdominal cavity. It may also be possible to implant a packed pellet of dye powder, similar to pharmacological manipulations using packed pellets of steroid hormone (e.g., Pradhan et al., 2014). The efficacy of the gelatin or pellet to dye eggs (a single clutch or multiple clutches) *in vivo* has yet to be tested.

We hope that as a result of this successful demonstration of egg marking in female *L. dalli* that others explore whether this method is useful in their research. While our primary goal was to estimate female fitness, we foresee additional applications, for example, in the study of development. To implement this method, we suggest piloting multiple colors of dye because both species and experimental context could affect the efficacy of the dye and/or confounds of specific dye colors. For example, although *L. dalli* females tolerated injections of orange dye well, eggs that were dyed orange could not be distinguished from the natural orange color of *L. dalli* eggs. Beyond the color of the egg, the external coloration of the female must also be considered because injecting dye temporarily colors the entire fish. This could affect social or reproductive behavior, or predation for field studies. Diluting the dye or waiting for the color to fade sufficiently before (re)placing the focal individual into the experimental context can address this issue. Interestingly, rose Bengal (red dye no. 105) had a similar effect on mortality in another goby species (Okuda et al., 2002) as the red dye (no. 40) used in this experiment, suggesting that red dyes may not be useful, in general, for these purposes.

Overall, this simple, inexpensive, and effective method for marking eggs makes it possible to quantify female reproductive success in polygynous fishes and other external fertilizers. Estimating fitness in more contexts and species will advance our understanding of evolution and the mechanisms underlying reproduction, development, and behavior.

## ACKNOWLEDGEMENTS

We thank A. Martinelli, A. Thomas, M. Williams, and L. Rogers for assistance with behavioral observations, quantifying reproduction, and fish maintenance and care. We thank the staff at the Wrigley Institute for Environmental Studies for technical support. This work was supported by grants from National Science Foundation to MSG (IOB-0548567), and National Science Foundation Doctoral Dissertation Improvement Grant (1311303), Sigma Xi, Honeycutt Fellowship, and Georgia State University Dissertation Grant, Brains & Behavior Program, and Neuroscience Institute to TKSL.

